# Quantifying the Cooperativity of Backbone Hydrogen Bonding

**DOI:** 10.1101/2025.03.29.646135

**Authors:** You Xu, Jing Huang

## Abstract

The hydrogen bonds (H-bonds) between backbone amide and carbonyl groups are fundamental to the stability, structure, and dynamics of proteins. A key feature of such hydrogen bonding interactions is that multiple H-bonds can enhance each other when aligned, as such in the *α* -helix or *β* -sheet secondary structures. To better understand this cooperative effect, we propose a new physical quantity to evaluate the cooperativity of intermolecular interactions. Using H-bond aligned N-methylacetamide molecules as the model system, we assess the cooperativity of protein backbone hydrogen bonds using quantum chemistry (QM) calculations at the MP2/aug-cc-pVTZ level, revealing cooperative energies ranging from 2 to 4.3 kcal/mol. A set of protein force fields was benchmarked against QM results. While the additive force field yielded null cooperativity, polarizable force fields, including the Drude and AMOEBA protein force fields, have been found to reproduce the trend of QM results, albeit with smaller magnitude. This work demonstrates the theoretical utility of the proposed formula for quantifying cooperativity and its relevance in force field parameterization. Incorporating cooperative energy into polarizable models presents a pathway to achieving more accurate simulations of biomolecular systems.

## 1 INTRODUCTION

Non-covalent interactions are essential for the structure and dynamics of biological macromolecules. When multiple molecules aggregate, their interactions can enhance each other due to the many-body effect. This is because the attractive contributions from dispersion and induction cooperatively increase with the number of molecules. This cooperative effect plays a vital role in the thermodynamics and kinetics of biomolecular processes such as folding, allostery, polymerization, and aggregation.[1, 2, 3, 4] Among non-covalent interactions, hydrogen bonds (H-bonds) are particularly notable for their cooperativity, as evidenced in the formation of secondary structures. For instance, H-bonds stabilize each other during the extension of *α* -helices and the accumulation of *β* -strands.[5, 6] These weak, cooperative and long-range interactions can be crucial for protein dynamics and functions.

To capture the cooperative effects of hydrogen bonding, computational chemistry has devoted significant efforts since its inception. Early quantum mechanical (QM) calculations employed Hartree-Fock (HF) methods combined with scaling factors, while modern density functional theory (DFT) methods offer greater feasibility. Despite advancements, precisely quantifying cooperativity using a defined formula remains a complex and debated issue in computational chemistry.[1] A pioneering study using a three-water system introduced a widely accepted formula for cooperativity in an *n*-molecule system[7], expressed as the difference between total energy and the sum of individual and pairwise interaction energies:

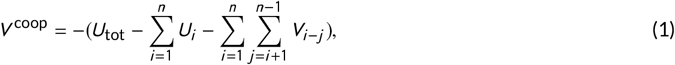

where *U*_*i*_ represents the internal energy of molecule *i*, and *V*_*i* −*j*_ denotes the pairwise interaction energy between molecules *i* and *j*. The minus sign ensures that attractive energy is represented as a positive value. This formula has been applied to many complex systems, including poly-water clusters,[8] protein building blocks such as N-methylacetamide (NMA),[9, 10] formamide,[11, 12] and *N* -methylformamide chains.[13] In these systems, cooperative energy typically accounts for 14-20% of the total interaction energy. Similarly in extendable *α* -helix contexts, the cooperativity contributes 2.6-3.2 kcal/mol for the first H-bond.[14, 15]

However, previous calculations did not fully isolate the cooperative effect on specific H-bonds while explicitly considering the geometric and basis set effects. Furthermore, cooperative energy arises from changes in the chemical environment, encompassing both intra- and intermolecular interactions.[16, 17] To properly account for geometric relaxation or deformation, energy calculations must employ identical molecular geometries across all energy terms in Eq. 1, using a consistent basis set to eliminate the basis set superposition error (BSSE).[18, 19]

Accurately capturing cooperative energy is also critical for developing transferable and accurate molecular mechanics (MM) force fields (FFs). In additive force fields, the absence of induction effects in backbone H-bonds is typically compensated by empirical scaling factors for dipole moments and corrections for backbone torsions. For example, *α* -helix formation simulated with the CHARMM and Drude force fields highlights this limitation.[20] Despite that, CHARMM FF is well-suited for analyzing conformational mechanisms of complex biomolecules, including membrane proteins [21, 22]. However, the intrinsic properties dominated by cooperativity, such as stabilized H-bonds in *α* -helices[14] or *β* -sheet aggregation in amyloid fibrils,[23][24] require explicit modeling of many-body effects. These effects, arising from the response of molecular charge redistribution, are a hallmark of polarizable force fields and must be rigorously evaluated during parameterization. Compared to additive FFs, the Drude model exhibits superior thermodynamic accuracy in describing metal-protein interactions [25] and buried ions [26], while AMOEBA excels in predicting hydrogen bonds and local p*K*_*a*_ shifts in response to electric fields in proteins. [27, 28]

In this work, we revisit the challenge of quantifying cooperativity using NMA as a model system. We propose an updated formula that isolates specific interactions while incorporating counterpoise (CP) corrections for BSSE and geometric relaxation. High-level *ab initio* calculations were employed to systematically examine the effects of NMA addition along different directions of a target dimer. The results are compared with previous studies to evaluate cooperativity, showing both similarities and differences. Furthermore, we assess the ability of CHARMM36, Drude (2013 and 2019), and AMOEBA (09 and newly-parametrized) force fields, providing insights into their strengths and limitations in modeling cooperative effects.

## 2 METHODS

### 2.1 Theoritical framework of quantifying cooperativity

In this section, we propose the equation for the cooperativity of interactions between two specific molecules. For a two-molecule system, *U*_*AB*_ represents the total energy, while *U*_*A*_ and *U*_*B*_ denote the internal energy of molecules *A* and *B*, respectively. The interaction energy *V*_*A*−*B*_ between these molecules is calculated as the difference between the total energy and the individual components:

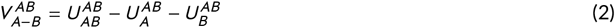

The subscript indicates the component of energy being considered, and the superscript *AB* denotes that the optimized geometry and the basis sets used for energy calculation are taken from the complex of molecules *A* and *B*. When a third molecule (denoted as *C*) is introduced, the total energy of the system becomes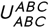, and the interaction energy between *A* and *B* is now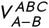. Due to many-body effects, a difference arises between 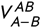 and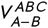, representing the cooperativity of the interaction energy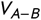:

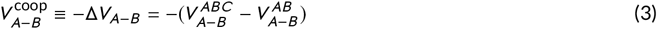

The interaction energy between *A* and *B* in the three-molecule system can be expressed as the difference between the total energy and all other internal and interaction energies:

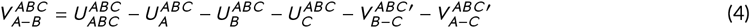

where

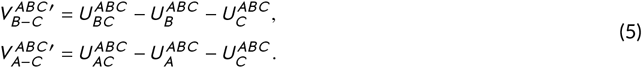

Substituting 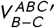 and 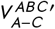 into eq. 4, and then substituting 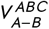 and 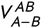 into eq. 3, the equation is rearranged as

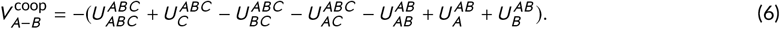

A positive 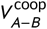 indicates that the addition of a molecule strengthens the interaction, and *vice versa*. Eq. 6 is the form of the cooperativity calculation proposed in this paper. It requires internal energy components from two sets of geometries. The first four terms are computed using the optimized geometry and complete basis set of the three-molecule system, whereas the remaining three terms are obtained from the tow-molecule system. This approach focuses on the many-body effect on the specific interaction between molecular bodies *A* and *B*. It can be generalized to any multi-molecule system in which body *C* represents the environment that influences the target interaction *V*_*A*−*B*_. All energy terms of a system are calculated using the identical geometry and basis set group, to include the effects of molecule complexation. Eq. 6 is therefore theoretically strict in accounting for molecular interactions.

It is important to note that eq. 4 assumes that the total energy of the system can be expressed as the sum of individual internal energies and pairwise interaction energies:

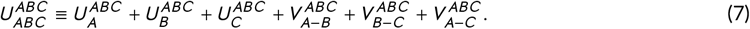

All energy components are derived from the same optimized geometry of the *ABC* complex. However, it is not possible to solve the terms of the interaction energy *V* analytically. The terms *V* calculated using eq. 5 accounts for geometric arrangements and consistent basis sets, but the effects of induced dipole moments caused by the electric field from absent molecules are still neglected. To fully resolve the interaction energy terms, a contribution of the induced dipole Δ*V*_*ϕ*_ must be included, ensuring that the sum of energy terms matches the total energy:

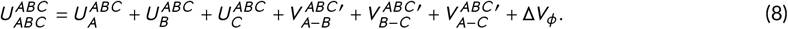

In eq 4, since the target interaction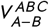 is calculated as the difference between the total energy and other pairwise interactions, the effect of induced dipole moment is inherently included:

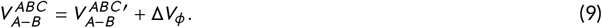

The term Δ*V*_*ϕ*_ contains the contributions of induction related to molecular bodies *A* and *B*, but not those within *C*. Thus, the cooperativity calculated using eq. 6 accounts for geometric rearrangements, counterpoise corrections of BSSE, and contributions of induced dipole moments.

### 2.2 Model systems and computational details

The molecular models are homogeneous NMA polymers in which NMAs were connected via hydrogen bonds (H-bonds) in a head-to-tail arrangement. The building block NMA dimer was organized in parallel configuration where the carbonyl oxygen of molecule *A* (O_*A*_) forms an H-bond with the amide nitrogen of molecule *B* (N_*B*_). In the *syn* conformation, atoms N_*A*_ and N_*B*_ are positioned on the same side of the C_*A*_-O_*A*_ axis, *i*.*e*., dihedral torsion ϕ_1_ (N_*A*_-C_*A*_-O_*A*_-N_*B*_) is 0^°^, while in the *anti* conformation, N_*A*_ and N_*B*_ lie on opposite sides of the C_*A*_-O_*A*_ axis, *i*.*e*., ϕ_1_ is 180^°^ (Figure 1A). Polymers composed of identical dimer conformations adopt an arc pattern, while polymers formed by alternating *syn* and *anti* dimers adopt a linear pattern (Figure 1B, C).

**FIGURE 1:**
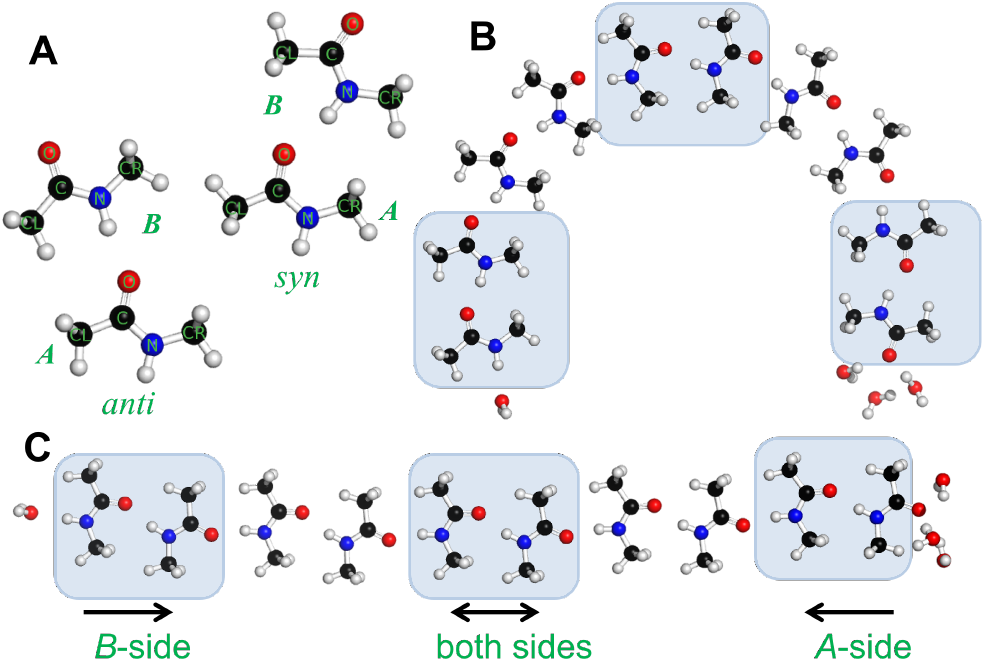
The chemical diagram of hydrogen-bond-aligned NMA models. The geometries are optimized in *ω*B97XD/cc-pVTZ level. Carbon atoms are shown in black, oxygen in red, nitrogen in blue, and hydrogen in white. (A) Two conformations of NMA dimerization, *syn* and *anti*, with labels indicating the molecules *A* and *B*. (B) An arc decamer exapmle consists exclusively of *syn* dimer conformations. (C) A linear decamer example consists of alternating *syn* and *anti* dimer conformations. Note the maximum of linear polymer is dodecamer. The cooperative energy is calculated for the first, middle, and last dimers of each polymer (highlighted with transparent boxes) to evaluate the effect of extending NMAs on *B* -side, both sides and *A*-side, respectively. In both (B) and (C), water molecules capping the terminal H-bond sites are illustrated, representing the case of 4-wat system.

All polymers were built starting from a *syn* dimer, and extended up to 10 NMAs in arc chains and 12 NMAs linear chains. The cooperativity was evaluated for the H-bonds that sit in the head, middle and tail of each polymer (Figure 1C), to evaluate the effect of additional NMA binding to molecule *B* (*B* -side), both molecules, and molecule *A* (*A*-side), respectively. The solvent effect was examined using the identical NMA polymers but the exposed amide and carbonyl groups of the terminal NMAs were capped with four water molecules: one was placed on the amide side to form a single H-bond, and three were positioned on the carbonyl side to form an H-bond network (Figure 1B, C). The polymers with or without solvent molecules were denoted as 4-wat and 0-wat systems, respectively.

All quantum mechanical (QM) calculations were performed in vacuum. Initial molecular conformations were built using the CHARMM program [29], with the initial O_*A*_-N_*B*_ distance setting to be 2.83 Å. Molecular geometries were optimized using Gaussian16 at the *ω*B97XD/cc-pVTZ level. To simplify structural diversity, the torsions *ϕ*_1_ and *ϕ*_2_ (C_*A*_-O_*A*_-N_*B*_ -C_*B*_) were constrained to ensure all NMA molecules in the plane. Single-point energy calculations were performed using Psi4 v1.15[30] at the RI-MP2/aug-cc-pVTZ level and the spin-component scaled MP2 (SCS-MP2) energy was adopted. Energy components were obtained on the individual geometries belonging to one complex while setting the remaining to be ghost atoms, enabling CP corrections for basis set superposition errors.

For molecular mechanics (MM) calculations, initial conformations were taken from the QM-optimized geometries. The same torsion constraints were used in MM minimization. The NMA parameters of three polarizable force fields, Drude-2013 (D13)[31, 32], Drude-2019 (D19)[33], AMOEBA09 (A09)[34, 35], and the up-to-date AMOEBA version parametrized using Poltype 2 (AMOEBA_Poltype2, Ap2)[36]; and one additive force field, CHARMM36 (C36)[37, 38], were employed. The Tinker package v8.6 [39] was used for AMOEBA09-related calculations, while CHARMM was used in calculations of the other force fields.

## 3 RESULTS

### 3.1 Cooperative energy in H-bonds

The NMA polymers were optimized using DFT and their energies were calculated using the MP2 method with a larger basis set. The interaction energy of an isolated dimer *AB* in the *syn* conformation is -6.19 kcal/mol, whereas it is slightly stronger at -6.75 kcal/mol in *anti*. The cooperative energy was evaluated using eq. 6. In all polymer patterns, the addition of NMA molecules enhanced the interactions of the original NMA dimers.

Adding one NMA on either the *A*- or *B* -side produced the cooperativity of 1.4 kcal/mol for the *syn* dimer, whereas a slightly lower energy of 1.1 kcal/mol was observed on the *A*-side extension of the *anti* dimer (Figure 2A, B). The cooperative energy, 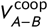, further increased and reached a plateau as the fifth NMA molecules was added for either conformation. The maximum cooperative energy was 2 kcal/mol for the *syn* conformation, with no significant difference between *A*- and *B* -side extensions. For *anti* conformation, it stabilized at 1.8 kcal/mol on the *A*-side elongation. Referred to eq 3, the lower cooperativity of *anti* correlates with the stronger isolated dimer interaction energy, so the difference of total interaction energy 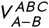 between two conformations is supposed to be tiny in a large NMA polymer.

**FIGURE 2:**
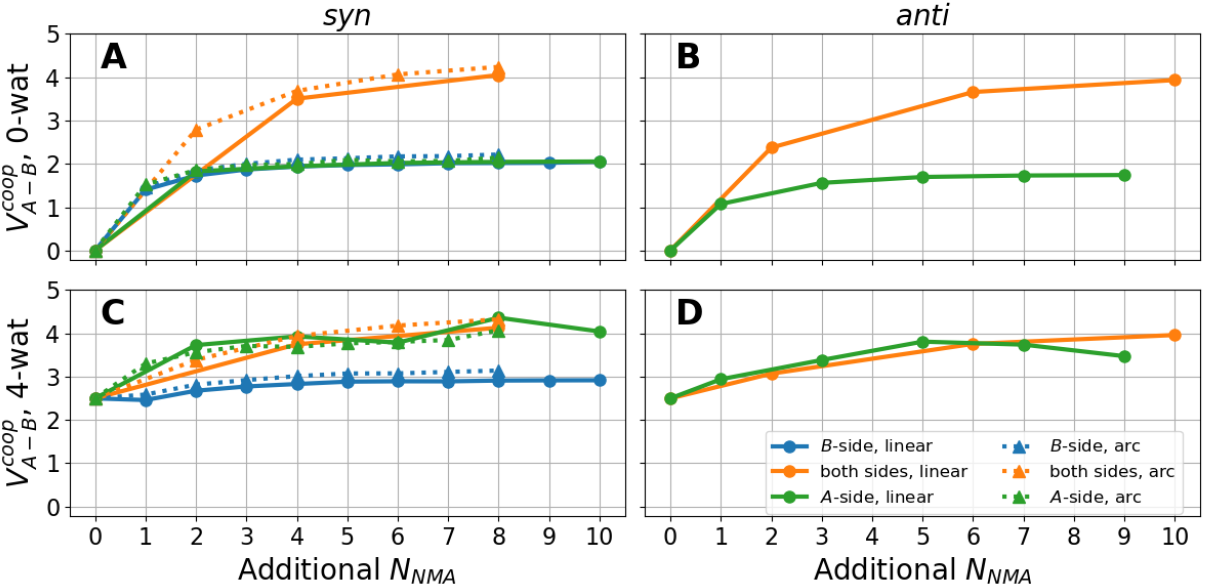
The QM-calculated cooperative energy between two NMAs. The *x* -axis represents the number of additional NMA molecules added to the target dimer, while the *y* -axis shows the cooperative energy in kcal/mol. Colors differentiate the extension directions of the NMA polymer: blue for the *B* -side, green for the *A*-side, and orange for both sides. Circles with solid lines represent data calculated for the line-pattern polymer, whereas triangles with dotted lines represent data for the arc-pattern polymer. (A) The *syn* dimer in the 0-wat system; (B) the *anti* dimer in the 0-wat system; (C) the *syn* dimer in the 4-wat system; (D) the *anti* dimer in the 4-wat system.

When NMA molecules were added to both sides of the dimer, the cooperative effect became more pronounced. The cooperative energy was 2.8 kcal/mol for the *syn* and 2.4 kcal/mol for the *anti* conformation when an NMA was added to both sides. As the polymer length increased, such energy exceeded 4 kcal/mol when a total of 10 NMA molecules were added on both sides (Figure 2A, B). Across all polymer lengths, the cooperativity contributed by each side was nearly identical, and the total effect on both sides was consistently double that of a single side. Additionally, elongation in the arc pattern produced slightly greater cooperative energy compared with the linear pattern. This is likely due to their closer spatial packing of NMA molecules, even though their dipole vectors are not as parallel as in linear chains.

To further assess the effect of solvent, the terminal donor and acceptor of all polymers were capped by water molecules. On the *A*-side, one water molecule forms a single H-bond with the amide group. On the *B* -side, three water molecules form two H-bonds with the carbonyl group and two additional H-bonds among themselves. Elongation of 4-wat systems therefore involves inserting NMAs between the corresponding terminal NMA and the water molecule(s). The cooperative energy from water molecules for the original dimer *AB* was 2.5 kcal/mol, which is comparable to adding an NMA to each side (Figure 2C). For the *syn* dimer, the cooperative energy further increased by approximately 0.5 kcal/mol, stabilizing at 3 kcal/mol. Compared to 2 kcal/mol in the corresponding 0-wat system, this 1 kcal/mol excess was attributed to the water molecule on the *A*-side. Adding the first NMA to the *B* -side of the *syn* dimer did not result in an additional increase in cooperativity, as this effectively replaced the H-bond between the molecule *B* and water molecules. This suggests that the cooperative effect contributed by NMA molecules is less significant than that of three water molecules. Comparing the 0-wat and 4-wat systems, the cooperative contribution of NMA lies between that of one and three water molecules.

For the *anti* dimer, the cooperativity from the *A*-side reached 3.8 kcal/mol, while for the *syn* dimer it increased to 4.3 kcal/mol. The more than 2 kcal/mol excess was attributed to the H-bond network formed by water molecules on the *B* -side. When NMAs were added to both sides of the dimer, the cooperative effect by water molecules was reduced, and the maximum cooperativity reached 4.3 kcal/mol, which was equivalent to the 0-wat system. Comparing the cooperativity in the 4-wat system, the influence of three water molecules was found to be even more significant than that of five linearly polymerized NMAs.

An unexpected decrease in cooperativity on *A*-side elongation was observed in the presence of water when the polymer length exceeded seven NMAs (Figure 2C, D). This decrease was primarily due to technical issues in the optimization process. As the system size increased, it became more challenging to maintain identical conformations for the newly added NMA units while enforcing all torsion constraints. Consequently, the imperfect conformations caused deviations in the cooperativity values. In summary, based on the calculations of all systems, the total interaction energies in *syn* and *anti* conformations are similar and the upper bound of cooperative energy in multi-NMA systems is estimated to be approximately 4.3 kcal/mol.

### 3.2 H-bond distance in NMA polymers

The hydrogen bond distances between O_*A*_ and N_*B*_ were 2.96 Å and 2.93 Å for *syn* and *anti* dimers, and their bond angles are 145^°^ and −150^°^ for C_*A*_-O_*A*_-N_*B*_, respectively. Upon the chain elongation, the angles showed minimal fluctuations around their original values, while the H-bond distances constantly decreased, correlating closely with the cooperative energy increment (Figure 3).

**FIGURE 3:**
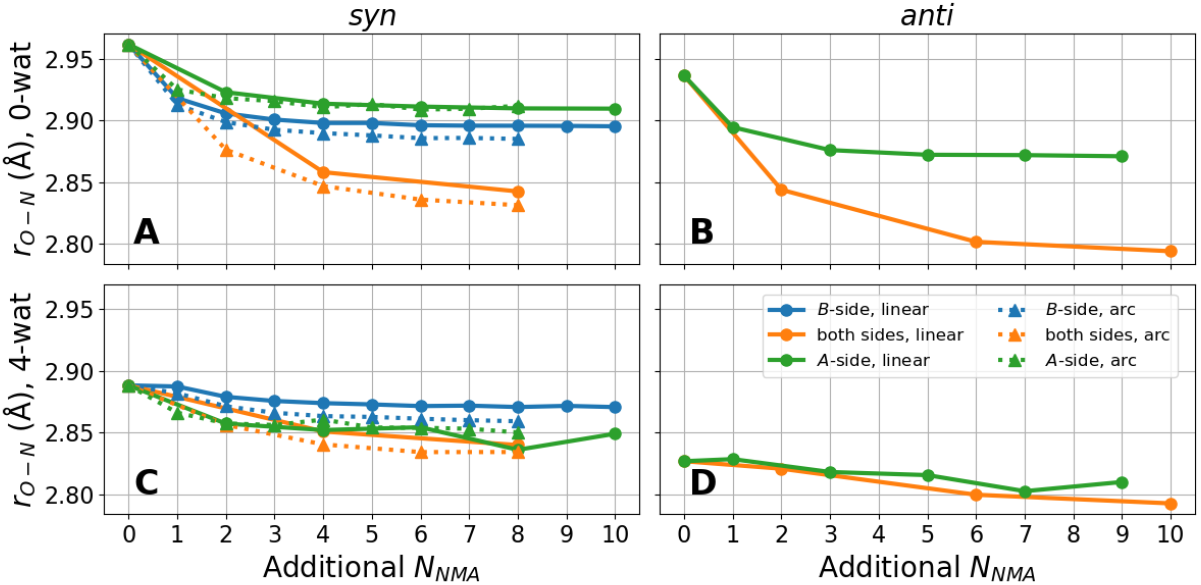
The QM-calculated H-bond distances between two NMAs. The *x* -axis represents the number of additional NMA molecules added to the target dimer, while the *y* -axis shows the distance between O_*A*_ and N_*B*_. Data organization and representation are identical to those in Figure 2.

The polymerization from *B* -side and both sides of the *syn* dimer reduced the H-bond distance to 2.88 Å and 2.83 Å, respectively. The reduction was slightly greater in arc chains compared to the linear ones. However, polymerization from the *A*-side resulted in no significant difference in H-bond distances between polymer shape. The *A*-side elongation led consistently less distance decrease than *B* -side in the 0-wat system, which means the H-bonds in the tail were slightly longer than that in the head of chains (Figure 3A). The H-bond distances of *anti* decreased more rapidly than *syn*, stabilizing at 2.87 Å and 2.79 Å as the NMA addition on the tail and middle H-bonds, respectively (Figure 3B), although the trends in cooperative energy increase were nearly identical for both conformations.

In the 4-wat system, the presence of water molecules significantly tightened the H-bonding interactions for the terminal dimers, particularly for dimers in the tail, which were capped with three water molecules. The H-bond distance reached 2.89 Å for the *syn* dimer and 2.83 Å for the *anti* dimer, matching the minimum distances observed in the 0-wat system. As a result, the contribution of additional NMAs replacing water molecules became small. This behavior of distance change upon NMA addition in different directions closely aligns with the trends in cooperative energy (Figure 3C). Consistent with the variations seen in cooperative energy, the deviations in H-bond distances observed during elongation from the *A*-side were attributed to poorly optimized geometries in longer chains.

### 3.3 Evaluation with additive force field

The cooperative energy calculated using SCS-MP2 at the aug-cc-pVTZ level was reproduced using force fields. The additive CHARMM36 FF was unable to reproduce cooperativity (Figure 4). The 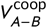 was nearly zero, showing only minor fluctuations in both 0-wat and 4-wat systems. Theoretically, in charge-fixed force fields, the terms 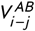 and 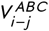 or 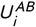 and 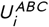 are almost identical. Without polarizability, there is no charge redistribution to account for environmental induction. As a result, the energetic differences between these terms arise from the conformational changes due to variations in H-bond distances, and are generally small for the NMA context. We emphasize that, although only C36 is tested here, any additive FF would exhibit nearly zero cooperativity due to their construct. The cooperativity defined in this study thus serves as an important example to demonstrate the inherit limitation of additive FF models.

**FIGURE 4:**
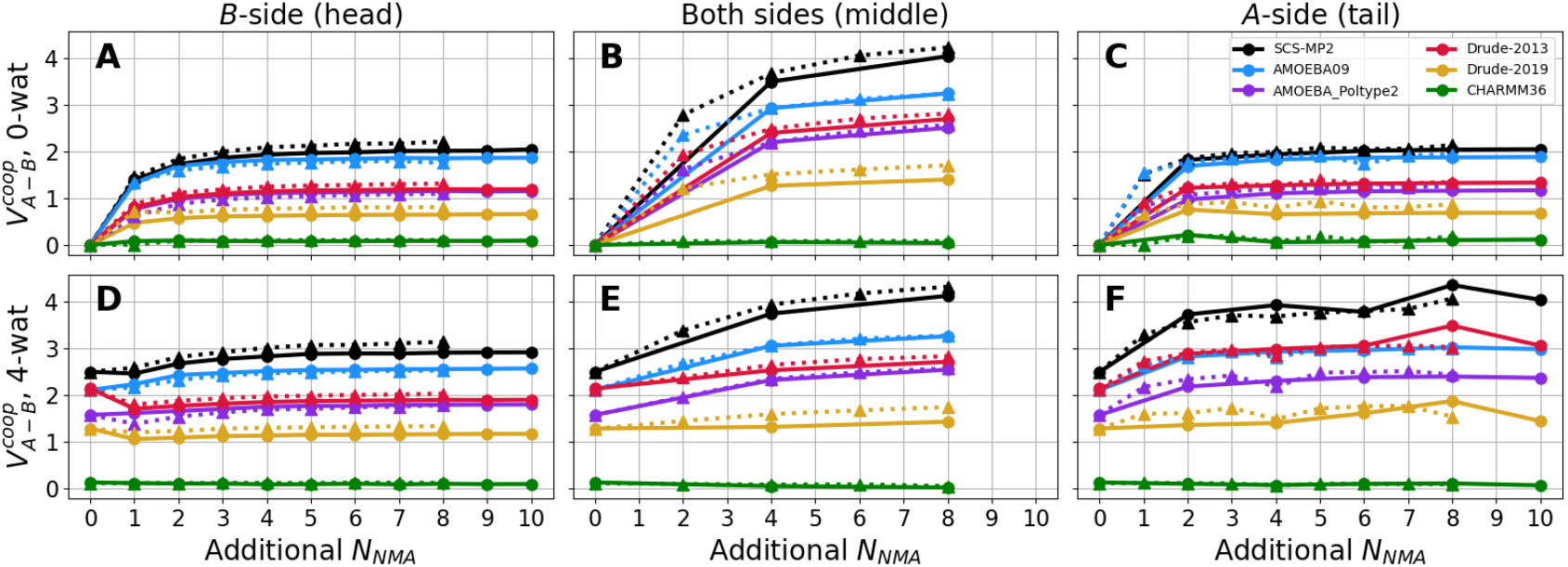
The comparison of cooperative energy for the *syn* conformation between QM and force field calculations. The *x* -axis represents the number of additional NMA molecules added to the target dimer, while the *y* -axis is the cooperative energy in kcal/mol. Colors differentiate the calculation methods: black for SCS-MP2, blue for AMOEBA09, violet for AMOEBA_Poltype2, red for Drude-2013, golden for Drude-2019, and green for CHARMM36m. Circles with solid lines represent data calculated for the linear polymer, whereas triangles with dotted lines represent for the arc polymer. (A) The *B* -side extension in the 0-wat system; (B) The both-side extension in the 0-wat system; (C) The *A*-side extension in the 0-wat system; (D) The *B* -side extension in the 4-wat system; (E) The both-side extension in the 4-wat system; (F) The *A*-side extension in the 4-wat system.

The H-bond distances in C36 were systematically shorter than those calculated by QM methods (Figure 5). Although the distances did not significantly change as the addition of NMA molecules, the overall trend still showed a slight decrease, particularly for the H-bond in the middle of the chain, which is consistent with QM results (Figure 5B, E). This suggests that the aggregation of NMA molecules enhances the long-range attractive interactions, thus compacting the polymers, despite the lack of explicit polarization effects in the additive force field.

**FIGURE 5:**
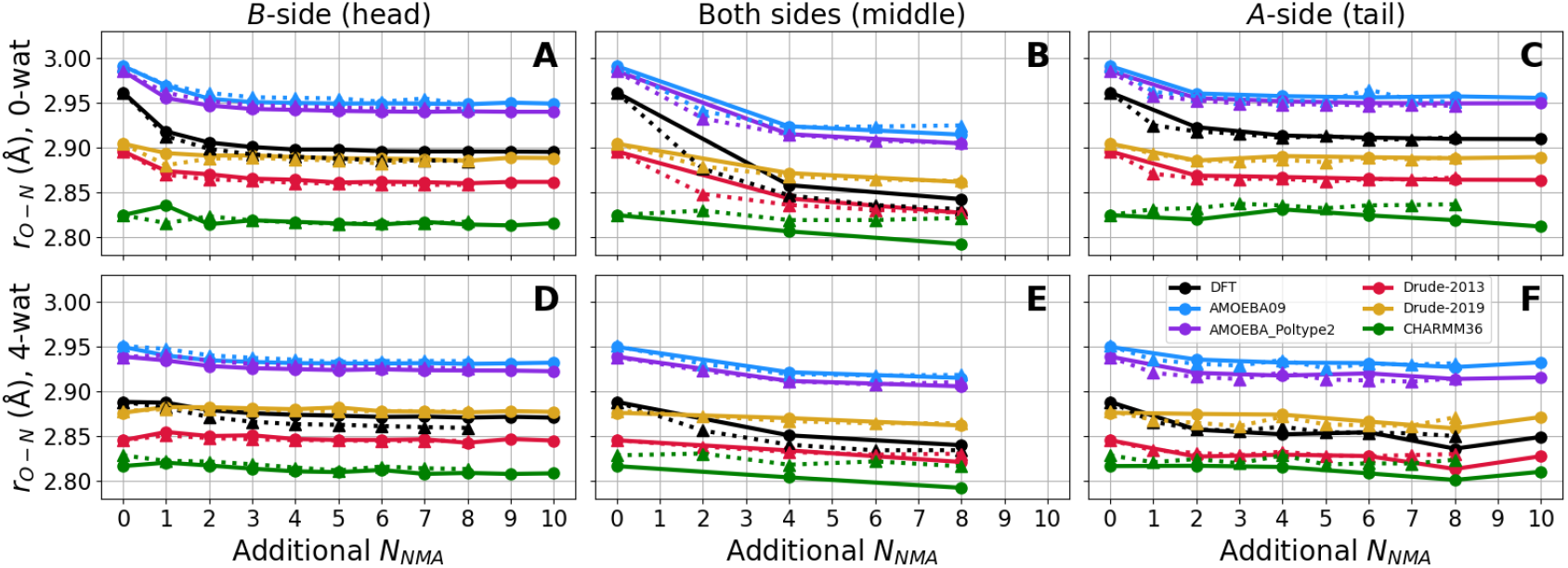
The comparison of H-bond distances for the *syn* conformation between QM and force field calculations. The *x* -axis represents the number of additional NMA molecules added to the target dimer, while the *y* -axis shows the distance between O_*A*_ and N_*B*_. Data organization and representation are identical to those in Figure 4.

### 3.4 Evaluation with polarizable force fields

In contrast to C36, polarizable force fields AMOEBA and Drude demonstrated the correct trends of cooperativity increment along with chain elongation. However, they systematically underestimated the magnitude, with deviations ranging from 10% to 60% (Figure 4A, B, C). Among them, AMOEBA09 achieved the best performance, particularly for elongation on the single side in the 0-wat system, where its results closely aligned with QM calculations. Nonetheless, its underestimation increased to 20% for interactions in the middle H-bomds. Drude-2013 and AMOEBA_Poltype2 underestimated cooperativity by approximately 40% across all elongation directions, while Drude-2019 showed the largest underestimation at 60%.

For a single H-bond in a dimer, the cooperative energy predicted by A09 and D13 was closer to the QM data, whereas Ap2 and D19 showed a certain degree of underestimation (Figure 4D). In the presence of water molecules, the overall trends observed in force field calculations remained consistent with QM calculations. For the middle H-bond, the maximum cooperative energy was identical between the 0- and 4-wat systems, as the effect from water was largely replaced by NMAs. A notable observation in Drude force fields was the decrease in cooperativity when an NMA was inserted between molecule *B* and the water molecules (Figure 4D). As a result, the cooperativity of the head H-bond in decamers was weaker compared to the contribution of three water molecules, while the cooperativity of the tail H-bond improved significantly and closely aligned with QM results. This indicates that the cooperative energy introduced by water molecules was comparable to MP2 results, but the weaker polarizability of NMAs in the Drude model led to the overall deviation.

The H-bond distances calculated using polarizable force fields displayed slightly different trends compared to their performance in energy. For isolated dimers, both AMOEBA models closely aligned with QM results, while both Drude force fields underestimated the H-bond distance by 0.06 Å, although they performed better than C36 (Figure 5A, B, C). However, all polarizable force fields underestimated the reduction in H-bond distances in response to chain elongation, consistent with their underestimation of cooperative energy. In longer NMA polymers the minimum distances of Drude models approached QM values, while AMOEBA models deviated further. Ultimately, Drude-2019 reproduced QM distances most accurately. In the 4-wat systems, the Drude force fields, particularly D19, closely matched QM results, whereas A09 and Ap2 overestimated the distances by 0.08 Å.

Overall, all polarizable force fields reproduced cooperative energy trends though underestimated their magnitudes, in stark contrast to additive force fields. Among the tested force fields, AMOEBA09 performed best for cooperative energy, while AMOEBA_Poltype2 and Drude-2013 exhibited further underestimation. The Drude-2019 models demonstrated potential for further improvement in capturing cooperativity across all systems.

## 4 DISCUSSION

### 4.1 Comparison with other methods to quantify cooperativity

The quantification of cooperative energy has been widely reported to yield varying values across different model systems. In earlier studies, limitations in computational resources often restricted the energy terms to being calculated using individually optimized geometries. Deviations in geometry potentially introduced undesirable geometric relaxation effects, further compounded by contamination from basis set superposition error (BSSE). Differences also arise from the various formulations used to define cooperative energy.

The basic concept originates from Eq. 1, which has been applied to directly calculate the total cooperative energy of a system. For example, studies on N-methylformamide (NMF) chains in the presence of a solvent molecule have reported cooperative energy contributions of up to 17.5 kcal/mol for a pentamer, representing 19% of the total interaction energy. [13] Another systematic QM study on water polymerization applied strict CP corrections. [8] In this case, the cooperative energy for a linear water heptamer was calculated to be 26 kcal/mol, accounting for 16% of the total energy. This effect increased to 18.6% when the water pentamer was arranged in a ring configuration.

In many other cases including this study, systems are simplified to three-body models, where two bodies form the target interaction and the third represents the environment. Eq. 1 therefore can be reformulated as either Eq. 10a or more practically using internal energy as shown in Eq. 10b.

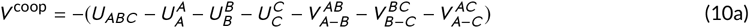

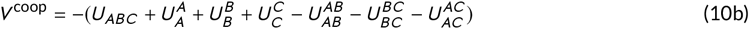

The difference between those formula and eq 6 is the superscripts, *i*.*e*., the consideration for the effect of geometry and basis set upon complexation. In a set of comprehensive studies for NMA structure and interactions, cooperative energy was calculated as the difference between trimer interaction energy and the interaction energies of two dimers.[9] This approach, based on Eq. 10a, omitted the −*V*_*A*−*C*_ term. As a result, the cooperative energy for adding an NMA molecule to the *A*-side of a dimer or three water molecules (one on the *A*-side, two on the *B* -side) enhanced the H-bond energy by 1.6 or 2.8 kcal/mol, respectively. [10] They are higher than our calculations which yield 1.4 kcal/mol (syn) or 1.1 kcal/mol (anti) for NMA and 2.4 kcal/mol for four-water addition (Fig. 2A, B). Although the differences in dimerization geometries (H-bonds along the C=O axis) and QM methods (HF/6-31G*) used in those studies might contribute the discrepancies, the omission of −*V*_*A*−*C*_ remains the primary cause of overestimation.

For some large systems, the cooperative energy has been quantified using the formulation equivalent to Eq. 11. Compared to Eq. 6, the term 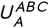 replacing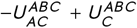, effectively adds the attractive term 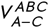 into the interaction energy. Consequently, elongation on both sides of the formamide dimer was reported to stabilize H-bond energy by 250%.[11]; the extension of the *α* -helix of glycine in one direction stabilized the H-bond to more than 3 kcal/mol[15]. In contrast, our study finds the stabilization on both sides of the NMA H-bond is only 67% and on one side is 2 kcal/mol, reflecting the difference of formulation rather than the varied H-bonds of molecular models.

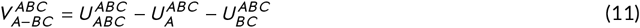

The cooperative effect on specific interactions between an alanine dipeptide and an extendable alanine helix was investigated to mimic the cooperative interactions of the first two H-bonds in a helix context. [14] The framework in this study is conceptually identical to this study, where the dipeptide was treated as *A*, the two alanines H-bonded to *A* as *B*, and the remaining alanines covalently bonded to *B* as *C*. The formula used can be written as Eq. 12, which is a form of Eq. 6 without superscripts, as the optimizations of whole-system geometry and BSSE corrections were not applied. Their reported cooperative energy of 2.6 kcal/mol closely aligns with our result of 2 kcal/mol for NMA addition on the *B* -side of linear NMA chains.

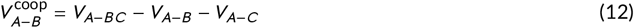

These findings suggest that H-bond enhancement in elongated helix contexts is approximately 30% for terminal H-bonds and 67% for middle H-bonds. This indicates that Eq. 6 is applicable even when *B* and *C* are covalently connected. Solving Eq. 6 for large systems can be computationally expensive, due to the need to evaluate terms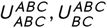, and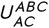, where *C* is usually the biggest part. However, advances in computational software and hardware have made it feasible to perform single-point energy calculations for sizable systems at very high theoretical levels. Advanced DFT methods combined with modern basis sets also make it practical to optimize the geometries of systems containing a few hundred atoms.

### 4.2 Significance in force field parametrization

Evaluating cooperative energy in MM calculations is crucial for validating force fields. In this study, NMA models from four force fields were assessed and compared with QM results. As the simplest building block of a protein backbone, NMAs can reflect the performance of these force fields in biomolecular modeling and simulations.

The NMA parameters in C36 were developed in CHARMM22 [37] and remain unchanged all the way towards the development of the current state-of-the-art CHARMM protein FF, CHARMM36m[40]. The NMA parameters in Drude-2013 were developed as amide model compounds for early Drude FFs [41], with their physical properties in condensed phase validated using Kirkwood-Buff analysis [42]. In contrast, the NMA parameters in Drude-2019 were directly derived from protein backbone parameters. Specifically, those parameters were initialized from Drude-2013 NMA, optimized using alanine peptides [31], and further refined for *β* -sheets and selected protein cases [33]. D19 remains the most up-to-date protein parameter set within the Drude force fields. On the other hand, the AMOEBA09 for small molecule was developed *ad hoc* as a independent proof of concept for AMOEBA parameterization [34]. The significantly improved multipole and bonded parameters for polypeptides were later incorporated into AMOEBA13 [43], which is now preferred for protein modeling. However, AMOEBA13 was not included in this study as it was not explicitly optimized for NMA. Instead, we included the parameters generated by Poltype 2, a recent automation protocol for AMOEBA [36], to represent the latest AMOEBA version for NMA. The water models remain identical across the versions of Drude[32] and AMOEBA[35] FFs used in this study. Consequently, NMA in C36 and D19, as backbone building blocks, represent the current protein force fields, while NMA in D13, A09, and Ap2 exist as model compounds and only partially reflect protein performance.

The performance of the force fields aligned with expectations: polarizable FFs reproduced cooperative energy to some extend, though both Drude and AMOEBA underestimated the magnitude. C36 reproduced the trend of compacted H-bond distances upon polymerization, but the effect was weaker than in polarizable force fields. The H-bond distance in C36 for isolated dimers was 0.14 Å shorter than QM geometries. However, this discrepancy decreased to 0.08 Å and 0.05 Å for terminal and middle dimers in longer chains, respectively. Compared to QM and experimental data, the underestimation of H-bond distance and orientation in additive force fields is a general issue [44]. Due to the lack of polarization and anisotropy, additive force fields often overestimate dimer H-bond interactions in a vacuum to remedy the cooperative effects in condensed phases. By combining these with correction terms, such as CMAP for backbone torsions[45, 46, 47], additive FFs can ultimately reproduce proper protein geometries in solution.

In the Drude models, D13 exhibited the best performance for monomer dipole moments in a vacuum and their induction in the presence of water molecules (Figure 6), consistent with previous validation studies [42]. However, the optimization of D19 parameters for polypeptide models led to a compromise in the accuracy of dipole moments, suggesting that the polarizability of NMA parameters was sacrificed during the adjustment for proteins. Compared with the differences in cooperative energy and dipole magnitude performance between D13 and D19, their induction responses remained similar. A09 also demonstrated insufficient induction for NMA, albeit stronger than the Drude models. Similarly, the updated Ap2 exhibited reduction in cooperative energy, suggesting that certain compromises were made in the development of AMOEBA datasets.

**FIGURE 6:**
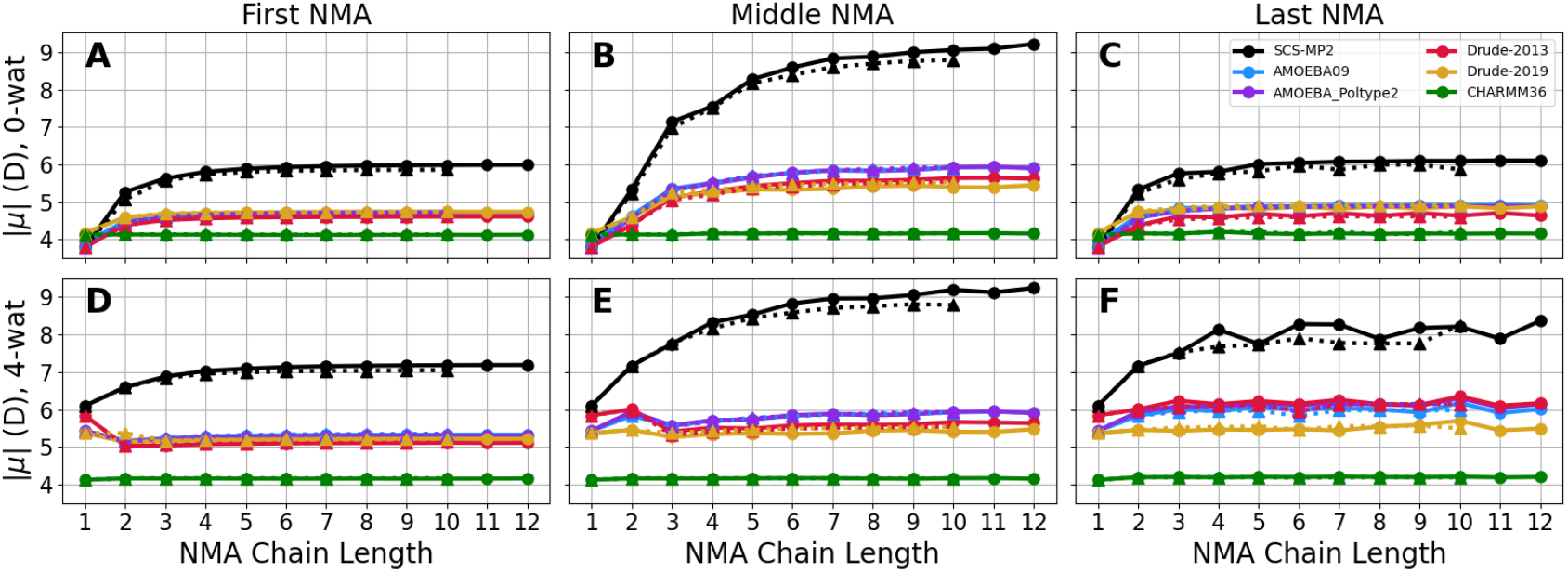
The dipole magnitude of NMA in different positions calculated using QM and force fields. The *x* -axis represents the chain length while the *y* -axis shows the molecular dipole moment in Debye. Shemoes of color and linestyle are identical to those in Figure 4. (A) The first NMA in the 0-wat system; (B) The middle NMA in the 0-wat system; (C) The last NMA in the 0-wat system; (D) The first NMA in the 4-wat system; (E) The middle NMA in the 4-wat system; (F) The last NMA in the 4-wat system.

The underestimation of cooperative energy in D19 is likely related to the reduced polarizability. In the Drude models, QM polarizability was scaled by 0.85 to better reproduce the dielectric constants of heteroaromatic compounds in aqueous phases [48] and this scheme was inherently adopted by protein FFs[31]. However, parameters fitted without scaling showed better performance for NMA dielectric constants, and cooperative energy calculated using Eq. 11 was superior to those with scaled parameters [41]. While reduced polarizability only slightly affects dipole moments, it has a significant impact on cooperative energy, influencing the entire system. Therefore the scaling scheme of polarizability in Drude protein FFs requires careful adjustment to balance the performance across compound classes.

Nonetheless, H-bond distances in D19 were more consistent with QM geometries in long chains compared to D13 and AMOEBA models. This is probably beneficial from that D19 parameters were optimized for polymer performance rather than single compounds. In summary, NMA in A09 performed best in cooperative energy and induced dipole moment but worse in H-bond distances. Ap2 had identical features in dipole magnitude and H-bind distance with A09, but further underestimated cooperative energy D13 excelled in water molecule cooperativity but underperformed in NMA binding. D19 showed reduced accuracy in cooperative energy and dipole moments due to scaled polarizability but reproduced H-bond distances better. As an additive FF, C36 could not explicitly describe cooperativity but achieved H-bond distances consistent with QM data for polymers.

We note that all four FF models included in this study are high-quality protein FFs, widely used in application studies of protein dynamics and function. The cooperative energy effectively differentiates the parameter performance in polarizability, even when dipole moments and interaction distances show minimal variation. This aspect may be overlooked in current biomolecular force field parametrizations. On the other hand, for comprehensive biomolecular parametrization protocols, accurately modeling bulk-phase properties may take precedence over strict agreement with gas-phase QM data. Therefore, underestimation of cooperativity can be justified if experimentally observed features are better reproduced. For any physics-based empirical potential, incorporating cooperative energy as a target during parameter optimization offers an additional functional for enhancing accuracy, though practical implementation requires further exploration.

Additionally, cooperativity could be used to validate and train machine learning potential (MLP) models.[49] One of the key challenges in MLP development is the capability to simulate larger and highly heterogeneous protein systems, where weak long-range interactions collectively act as the driving forces behind various processes [50, 51]. Using cooperativity data to complement existing training or validation datasets, and incorporating explicit cooperativity metrics into the loss function, may offer a means to enhance the long-range perception of MLPs in complex systems. To support these efforts, we provide QM-optimized structures and corresponding energy terms as Supporting Information. Table S1 and S2 include the energy terms calculated for arc and linear chains without water, respectively. Table S3 and S4 include the energy terms calculated for arc and linear chains with water molecules, respectively. Table S5 includes the energy terms for dimers. Those data offer benchmarks for force fields, whether MM- or ML-based, targeting NMA models by other researchers.

## 5 CONCLUSIONS

We proposed a formula for calculating the cooperative energy of non-bonded interactions. This approach treats any system as classical three-body interactions, focusing on the interaction between targets *A* and *B* while accounting for the cooperative effect induced by the environment *C*. Unlike previous methods, it incorporates the effects of geometric relaxation and induced dipole moments on the *A*-*B* interaction but excludes interactions with or within *C*. DFT calculations with a large basis set on NMA polymers revealed that the cooperative energy of hydrogen bonds ranges from 2 to 4.3 kcal/mol, with the energy in the middle of the chain being twice that of the terminal. While polarizable force fields successfully reproduced the cooperative effects, they underestimated the magnitude of the cooperative energy. In contrast, the additive force field failed to capture the cooperativity entirely. Our results highlight the theoretical utility of eq 6 for quantifying cooperativity in computational chemistry and provide insights for further refinement of FFs. This study underscores the importance of capturing polarizable features in FF development, in addition to fitting QM dipole moments, polarizabilities and geometries.

## Abbreviations

NMA: N-methylacetamide (NMA)
H-bond: hydrogen bond
QM: quantum mechanics
MM: molecular mechanics

## Acknowledgements

Y. X. thanks the discussion with Dr. Zilin Song. This work was supported by the “Pioneer” and “Leading Goose” R&D Program of Zhejiang (2023C03109, 2024SSYS0036), the National Natural Science Foundation of China (32171247, 21803057). We thank the Westlake University Supercomputer Center for computational resources and related assistance.

## Supporting Information

Supporting Information includes the QM optimized geometries of NMA polymers and the energy terms for cooperativity calculation used in this study.

