## Supplemental Materials for "Quantifying the Cooperativity of Backbone Hydrogen Bonding"

### Supporting Information for: Quantifying the Cooperativity of Backbone Hydrogen Bonding

You Xu[1,2,3] and Jing Huang*[1,2,3]

[1]Key Laboratory of Structural Biology of Zhejiang Province, School of Life Sciences, Westlake University, Hangzhou, Zhejiang 310024, China
[2]Westlake AI Therapeutics Laboratory, Westlake Laboratory of Life Sciences and Biomedicine, Hangzhou, Zhejiang 310024, China
[3]Institute of Biology, Westlake Institute for Advanced Study,Hangzhou, Zhejiang 310024, China

*No. 600 Dunyu Road, Xihu District, Hangzhou, Zhejiang, 310030, P.R. China;

Table S1: Energy terms of NMA chains in the 0-wat and arc systems. Within each chain, the molecules forming the first, middle, and last hydrogen bonds were designated as $A$ and $B$, respectively, while the remaining molecules were considered as $C$. Spin-component scaled MP2 energies were computed using Psi4 v1.15 at the RI-MP2/aug-cc-pVTZ level. All energy values are reported in Hartree.

| $N_{\mathrm{NMA}}$ | Position | $U_{ABC}^{ABC}$ | $U_{BC}^{ABC}$ | $U_{AC}^{ABC}$ | $U_{C}^{ABC}$ |
|---|---|---|---|---|---|
| 3 | First | -744.2146024776083095 | -496.1379027616911799 | -496.1274897465120830 | -248.0629801443959650 |
| | Middle | | -496.1379027616911799 | -496.1274897465120830 | -248.0629801443959650 |
| | Last | | -496.1384841575948599 | -496.1274897465120830 | -248.0636579977437179 |
| 4 | First | -992.2930361176936458 | -744.2152680192783691 | -744.2028199392291299 | -496.1378557174114121 |
| | Middle | | -744.2028199392291299 | -744.2027707742406619 | -496.1268607434228102 |
| | Last | | -744.2159285935802018 | -744.2027707742406619 | -496.1384834174691036 |
| 5 | First | -1240.3718955777048905 | -992.2936304407933221 | -992.2803728555120415 | -744.2151554808201581 |
| | Middle | | -992.2803728555120415 | -992.2783368965224327 | -744.2018679714710743 |
| | Last | | -992.2943698885388812 | -992.2804341715090004 | -744.2159105112272073 |
| 6 | First | -1488.4512407948798227 | -1240.3727081565546086 | -1240.3590666582706490 | -992.2937372950605095 |
| | Middle | | -1240.3562721574242005 | -1240.3563454614607053 | -992.2771095557551462 |
| | Last | | -1240.3731854835934882 | -1240.3593287470559972 | -992.2943126299804817 |
| 7 | First | -1736.5302253815702898 | -1488.4515494434608627 | -1488.4377304424745034 | -1240.3723078659375005 |
| | Middle | | -1488.4345961033718595 | -1488.4339561273302479 | -1240.3544225827399714 |
| | Last | | -1488.4522943243955524 | -1488.4379025695575365 | -1240.3731637624910036 |
| 8 | First | -1984.6098412840694891 | -1736.5310400150397072 | -1736.5171427363770817 | -1488.4516652109655297 |
| | Middle | | -1736.5128824066327979 | -1736.5129227304119013 | -1488.4323023199376621 |
| | Last | | -1736.5315332092991412 | -1736.5174359373120296 | -1488.4522667145463402 |
| 9 | First | -2232.6891708247567294 | -1984.6102892988583335 | -1984.5963119580096645 | -1736.5307635569852209 |
| | Middle | | -1984.5918503587408850 | -1984.5915891688512147 | -1736.5108062999506728 |
| | Last | | -1984.6108038570253029 | -1984.5966788069288214 | -1736.5314672263775719 |
| 10 | First | -2480.7683802017863854 | -2232.6894096740384157 | -2232.6754031235382172 | -1984.6098244767927099 |
| | Middle | | -2232.6704865372521454 | -2232.6705233007737661 | -1984.5892440838779294 |
| | Last | | -2232.6901801952080859 | -2232.6756648646423855 | -1984.6107417848400019 |

Table S2: Energy terms of NMA chains in the 0-wat and linear systems. Within each chain, the molecules forming the first, middle, and last hydrogen bonds were designated as $A$ and $B$, respectively, while the remaining molecules were considered as $C$. Spin-component scaled MP2 energies were computed using Psi4 v1.15 at the RI-MP2/aug-cc-pVTZ level. All energy values are reported in Hartree.

| $N_{\mathrm{NMA}}$ | Position | $U_{ABC}^{ABC}$ | $U_{BC}^{ABC}$ | $U_{AC}^{ABC}$ | $U_{C}^{ABC}$ |
|---|---|---|---|---|---|
| 3 | First | -744.2147753996437132 | -496.1381727094901066 | -496.1276509226831877 | -248.0631452951174083 |
| | Middle | | -496.1381727094901066 | -496.1276509226831877 | -248.0631452951174083 |
| | Last | | -496.1383037348986136 | -496.1276509226831877 | -248.0636507574695884 |
| 4 | First | -992.2929871103260666 | -744.2153521193465622 | -744.2026172006958404 | -496.1376021964434244 |
| | Middle | | -744.2026172006958404 | -744.2026793283313282 | -496.1268512765350351 |
| | Last | | -744.2158247509466946 | -744.2026793283313282 | -496.1382888486215279 |
| 5 | First | -1240.3721063296125067 | -992.2939884468179343 | -992.2804093681833137 | -744.2151328050438224 |
| | Middle | | -992.2804093681833137 | -992.2786289781505502 | -744.2021836985368282 |
| | Last | | -992.2941850715612873 | -992.2804778927709322 | -744.2158027554053206 |
| 6 | First | -1488.4511762432312025 | -1240.3728379464703266 | -1240.3588924527778090 | -992.2935031419311827 |
| | Middle | | -1240.3563921952602414 | -1240.3560362801893007 | -992.2766972037070445 |
| | Last | | -1240.3730795589171976 | -1240.3592512913796782 | -992.2941138542536237 |
| 7 | First | -1736.5303895401011687 | -1488.4518990557066900 | -1488.4378058092438550 | -1240.3723218991790418 |
| | Middle | | -1488.4349329537394624 | -1488.4338489385922912 | -1240.3541515052638715 |
| | Last | | -1488.4521025175906743 | -1488.4378686779011787 | -1240.3730457195006238 |
| 8 | First | -1984.6097740066727511 | -1736.5311973844643489 | -1736.5170211401034521 | -1488.4514747196719782 |
| | Middle | | -1736.5125392062320770 | -1736.5130218190206506 | -1488.4323755982395596 |
| | Last | | -1736.5313998050842201 | -1736.5173630787651291 | -1488.4520701441340407 |
| 9 | First | -2232.6890477191773243 | -1984.6104021684222971 | -1984.5961712400023771 | -1736.5305990017111526 |
| | Middle | | -1984.5914826454720696 | -1984.5916514389359691 | -1736.5108120581508047 |
| | Last | | -1984.6106336301440933 | -1984.5962398684200707 | -1736.5313436981562063 |
| 10 | First | -2480.7685147778861392 | -2232.6898398353478115 | -2232.6755662423843205 | -1984.6099711946440038 |
| | Middle | | -2232.6708043490434648 | -2232.6703376759369348 | -1984.5889431115112984 |
| | Last | | -2232.6900487530278951 | -2232.6759134170911238 | -1984.6105694283080538 |
| 11 | First | -2728.8477886164641859 | -2480.7690940680640779 | -2480.7547645089234720 | -2232.6891568547125644 |
| | Middle | | -2480.7498799817690269 | -2480.7492914672643565 | -2232.6677785366541684 |
| | Last | | -2480.7693177537435076 | -2480.7548745293611319 | -2232.6899393366566073 |
| 12 | First | -2976.9272098625324361 | -2728.8484597571414270 | -2728.8341588871776366 | -2480.7685375836813364 |
| | Middle | | -2728.8285226606017204 | -2728.8290128914181878 | -2480.7473543608844011 |
| | Last | | -2728.8487155396405797 | -2728.8345683069801453 | -2480.7692089473471242 |

Table S3: Energy terms of NMA chains in the 4-wat and arc systems. Within each chain, the molecules forming the first, middle, and last hydrogen bonds were designated as $A$ and $B$, respectively, while the remaining molecules were considered as $C$. Spin-component scaled MP2 energies were computed using Psi4 v1.15 at the RI-MP2/aug-cc-pVTZ level. All energy values are reported in Hartree.

| $N_{\mathrm{NMA}}$ | Position | $U_{ABC}^{ABC}$ | $U_{BC}^{ABC}$ | $U_{AC}^{ABC}$ | $U_{C}^{ABC}$ |
|---|---|---|---|---|---|
| 2 | First | -801.4913228369969147 | -553.4064781456894480 | -553.3912979539384196 | -305.3202938339098296 |
| | Middle | | -553.4064781456894480 | -553.3912979539384196 | -305.3202938339098296 |
| | Last | | -553.3912979539384196 | -553.4064781456894480 | -305.3202938339098296 |
| 3 | First | -1049.5698355159811399 | -801.4837059491393347 | -801.4780930103148648 | -553.4059445931412711 |
| | Middle | | -801.4837059491393347 | -801.4780930103148648 | -553.4059445931412711 |
| | Last | | -801.4678360665074024 | -801.4780930103148648 | -553.3912120957177194 |
| 4 | First | -1297.6487467971264778 | -1049.5618388204359235 | -1049.5556113805071163 | -801.4830496875837298 |
| | Middle | | -1049.5556113805071163 | -1049.5549026487763058 | -801.4770235746534581 |
| | Last | | -1049.5459730609966300 | -1049.5549026487763058 | -801.4676649729813107 |
| 5 | First | -1545.7278378474870806 | -1297.6405431058508384 | -1297.6338574791996052 | -1049.5610764414186633 |
| | Middle | | -1297.6338574791996052 | -1297.6326208731322822 | -1049.5542818058008834 |
| | Last | | -1297.6246258447547461 | -1297.6331169186398711 | -1049.5456751858532698 |
| 6 | First | -1793.8069657145383644 | -1545.7193776386891386 | -1545.7124806874915066 | -1297.6395513136499176 |
| | Middle | | -1545.7107324170483480 | -1545.7105925792789094 | -1297.6304954713521056 |
| | Last | | -1545.7050887791679088 | -1545.7111471680775594 | -1297.6250032675370676 |
| 7 | First | -2041.8863571571280318 | -1793.7986353423359560 | -1793.7916125459071282 | -1545.7186481262303914 |
| | Middle | | -1793.7897503029546442 | -1793.7893004631303029 | -1545.7090259185092691 |
| | Last | | -1793.7828853814462491 | -1793.7910012144570828 | -1545.7033932746076061 |
| 8 | First | -2289.9656966407542313 | -2041.8778733478120557 | -2041.8707796705368764 | -1793.7977171198399446 |
| | Middle | | -2041.8681919991640825 | -2041.8682687685304700 | -1793.7872828362308155 |
| | Last | | -2041.8621121595792829 | -2041.8702106088387609 | -1793.7825446220731465 |
| 9 | First | -2538.0452330724319836 | -2289.9572940121583997 | -2289.9502192476575146 | -2041.8770946385122897 |
| | Middle | | -2289.9474425264238562 | -2289.9472698982485781 | -2041.8661484842853042 |
| | Last | | -2289.9415113195082085 | -2289.9496014183605439 | -2041.8618667131690927 |
| 10 | First | -2786.1247535355887521 | -2538.0366269486557940 | -2538.0295073929919454 | -2289.9562451961301122 |
| | Middle | | -2538.0265252963490639 | -2538.0265878502109445 | -2289.9451078264551143 |
| | Last | | -2538.0189153778469517 | -2538.0277690945126778 | -2289.9382776116904097 |

Table S4: Energy terms of NMA chains in the 4-wat and linear systems. Within each chain, the molecules forming the first, middle, and last hydrogen bonds were designated as $A$ and $B$, respectively, while the remaining molecules were considered as $C$. Spin-component scaled MP2 energies were computed using Psi4 v1.15 at the RI-MP2/aug-cc-pVTZ level. All energy values are reported in Hartree.

| $N_{\mathrm{NMA}}$ | Position | $U_{ABC}^{ABC}$ | $U_{BC}^{ABC}$ | $U_{AC}^{ABC}$ | $U_{C}^{ABC}$ |
|---|---|---|---|---|---|
| 2 | First | -801.4913228536121323 | -553.4064781264482917 | -553.3912979897326068 | -305.3202938114745848 |
| | Middle | | -553.4064781264482917 | -553.3912979897326068 | -305.3202938114745848 |
| | Last | | -553.3912979897326068 | -553.4064781264482917 | -305.3202938114745848 |
| 3 | First | -1049.5697809804278222 | -801.4839043803311824 | -801.4777232986332365 | -553.4056257628727735 |
| | Middle | | -801.4839043803311824 | -801.4777232986332365 | -553.4056257628727735 |
| | Last | | -801.4677732490113158 | -801.4777232986332365 | -553.3911578322361038 |
| 4 | First | -1297.6486001278840376 | -1049.5618174257294868 | -1049.5550478487407418 | -801.4823870643999726 |
| | Middle | | -1049.5550478487407418 | -1049.5536193331824961 | -801.4757148715621042 |
| | Last | | -1049.5435423549010920 | -1049.5536193331824961 | -801.4643616481187109 |
| 5 | First | -1545.7278021542038005 | -1297.6406847662658492 | -1297.6337085886507339 | -1049.5608699854844872 |
| | Middle | | -1297.6337085886507339 | -1297.6327839338769081 | -1049.5544883153274895 |
| | Last | | -1297.6245427123942591 | -1297.6326806350912193 | -1049.5455689542684468 |
| 6 | First | -1793.8068619945790942 | -1545.7195187095874189 | -1545.7122749613470205 | -1297.6393010460615187 |
| | Middle | | -1545.7108360923934924 | -1545.7101885881490944 | -1297.6299929136080209 |
| | Last | | -1545.7011591467719427 | -1545.7104629242526244 | -1297.6208812994680102 |
| 7 | First | -2041.8857915174619393 | -1793.7983107841291712 | -1793.7909640339687485 | -1545.7179392043151438 |
| | Middle | | -1793.7893902901423644 | -1793.7884262819486594 | -1545.7080007644840407 |
| | Last | | -1793.7797526393642329 | -1793.7892815302918734 | -1545.7000677878570514 |
| 8 | First | -2289.9655827423534902 | -2041.8780355044439148 | -2041.8705732249220546 | -1793.7974955055924511 |
| | Middle | | -2041.8678293159107398 | -2041.8682822607527214 | -1793.7872706802072571 |
| | Last | | -2041.8618584187088345 | -2041.8700241748556437 | -1793.7821892302326887 |
| 9 | First | -2538.0441432264478863 | -2289.9565691067477928 | -2289.9491166854286348 | -2041.8760092436075411 |
| | Middle | | -2289.9461711902940806 | -2289.9463787857889656 | -2041.8652222257755966 |
| | Last | | -2289.9389596286609958 | -2289.9470171877510438 | -2041.8585485381067883 |
| 10 | First | -2786.1188946099709938 | -2538.0312451561135276 | -2538.0237391245727849 | -2289.9505858278389496 |
| | Middle | | -2538.0207962296130972 | -2538.0202254384603293 | -2289.9385667080832718 |
| | Last | | -2538.0113061786823891 | -2538.0209361842094040 | -2289.9301554617741203 |
| 11 | First | -3034.2035219910007982 | -2786.1158612284234550 | -2786.1083475300647478 | -2538.0351897883929269 |
| | Middle | | -2786.1053869577649493 | -2786.1048060353919027 | -2538.0231253582210229 |
| | Last | | -2786.1011168671270752 | -2786.1070188580833928 | -2538.0209033602068303 |
| 12 | First | -3282.2831068958748801 | -3034.1954290251096609 | -3034.1879037109733872 | -2786.1147322437482217 |
| | Middle | | -3034.1842426505982075 | -3034.1847230179678263 | -2786.1029272365321958 |
| | Last | | -3034.1769761568166359 | -3034.1860730388243610 | -2786.0962432879041444 |

Table S5: Energy terms of NMA dimer $AB$ in two conformations. Spin-component scaled MP2 energies were computed using Psi4 v1.15 at the RI-MP2/aug-cc-pVTZ level. All energy values are reported in Hartree.

| conformation | $U_{AB}^{AB}$ | $U_{A}^{AB}$ | $U_{B}^{AB}$ |
|---|---:|---:|---:|
| *syn* | -496.1357755926256345 | -248.0631137527353758 | -248.0627992554040873 |
| *anti* | -496.1373843165470134 | -248.0634597764712339 | -248.0631643312641188 |
